# Effect of a low water concentration in chloride, sodium and potassium on oocyte maturation, oocyte hydration, ovulation and egg quality in rainbow trout

**DOI:** 10.1101/2021.06.22.449391

**Authors:** Emilien Segret, Emilie Cardona, Sandrine Skiba-Cassy, Frédéric Cachelou, Julien Bobe

## Abstract

Water salinity is an important environmental factor known to have detrimental effects on salmonid reproduction, mostly when migrating female broodfish are held in sea water. In contrast, data obtained in freshwater are scarce and the impact of low water salinity, and more specifically of low water concentrations in sodium, chloride and potassium, during reproduction in freshwater is currently unknown. For this reason, and because ion and water fluxes are critical for the final steps of the female gamete formation, including oocyte hydration and ovulation, the aim of the present study was to investigate the impact of low salinity water on final oocyte maturation, ovulation and, ultimately, on egg quality, using rainbow trout as a physiological model and relevant aquaculture species.

Fish from the same commercial strain were raised either in a site characterized by low concentrations of Na+, K+, and Cl- ions in the water or in a closely located control site exhibiting higher concentration in these elements. Egg quality and duration of final oocyte maturation were investigated using innovative phenotyping tools such as automatic assessment of egg viability using the VisEgg system and non-invasive echograph-based monitoring of final oocyte maturation duration, respectively. Oocyte hydration during final oocyte maturation and after ovulation was also investigated. Finally, molecular phenotyping was performed using real-time PCR-based monitoring of several key players of final oocyte maturation and ovulation associated with ion and water transport, inflammation, proteolytic activity, and coagulation. Oocyte hydration and gene expression data were analyzed in the light of the duration of final oocyte maturation.

Here we show that low water salinity (i.e., low water concentration in chloride, sodium and potassium) negatively influences final oocyte maturation, ovulation and, ultimately, egg quality. Low water salinity triggered delayed ovulation and lower oocyte viability. When investigating the impact of low water salinity on final oocyte maturation duration, individuals presenting the most severe phenotypes exhibited impaired oocyte hydration and abnormally reduced gene expression levels of several key players of the ovulatory process. While the under expression of water (i.e., aquaporins) and ion (i.e., solute carriers) transporters is consistent with impaired oocyte hydration, our observations also indicate that the overall ovulatory gene expression program is disrupted. Our results raise the question of the mechanisms underlying the negative influence of low salinity water on the dynamics of the preovulatory phase, on the control of the oocyte homeostasis, including hydration, and on the overall success of the maturation-ovulation process.

**Highlights:**

- Low water salinity impairs final oocyte maturation and egg quality in rainbow trout
- Low water salinity induces delayed ovulation and impaired oocyte hydration
- Low water salinity induces a dysregulation of several key ovulatory genes
- Monitoring of final oocyte maturation can be performed using ultrasound staging

## 1 Introduction

Fish egg quality is defined as the egg ability to be fertilized and to subsequently develop into a normal embryo (Bobe and Labbé, 2010). Egg quality can be highly variable in fish under both natural and aquaculture conditions. In most species, a lack of knowledge of mechanisms influencing egg quality is a limiting factor for the mass production of fry (Migaud et al. 2013). Many external factors such as broodstock nutrition, temperature and photoperiod, to name just a few, are known to have a major impact on egg quality in various fish species (Aegerter and Jalabert, 2004; Bonnet et al., 2007; Carrillo et al., 1989; Izquierdo et al., 2001; Pankhurst et al., 1996). In contrast, other factors such as composition and physico-chemical parameters of the water have received far less attention (Brooks et al., 1997; Bobe and Labbe, 2010). Ionic composition of the water is however likely to influence the final steps of oogenesis (i.e., final oocyte maturation - FOM) given the importance of water exchange during this critical period (Dolomatov, 2012). Among mineral elements, chloride, sodium and potassium are of specific interest because (i) they are the most abundant electrolytes in the body of living organisms and (ii) they serve a vital function in controlling osmotic pressure and acid-base equilibrium (Lall, 2003). It is commonly accepted that deficiencies in chloride, sodium and potassium are difficult to trigger because fish derive these mineral elements from surrounding water (Lall, 2003). To our knowledge, the impact of a low ionic composition in these three key mineral elements on female reproduction was never investigated.

Several concomitant, yet distinct, events take place in the full-grown ovarian follicle during FOM. The follicle-enclosed oocyte progressively acquires the ability to resume meiosis whereas the follicle is preparing the release of oocyte from surrounding somatic layers at ovulation (Grier et al., 2007; Lubzens et al., 2010). In fish, as in other vertebrates, the mechanisms involved in the ovulatory process are described as inflammatory-like (Espey, 1994; Thibault and Levasseur, 1988). In contrast to meiosis resumption and ovulation that are common to all vertebrates, FOM in fish is associated with a significant hydration of the oocyte. This phenomenon results in a dramatic increase of oocyte volume in marine species and is triggered by water entry due to the increase in free amino acid content resulting from yolk protein processing (Finn et al., 2002). In the saltmarsh species, Fundulus heteroclitus, the influx in potassium and sodium is a major cause of the uptake of osmotically obligated water and subsequent volume increase experienced by maturing oocytes (Wallace et al., 1992). In fresh water species, a more limited, yet significant, increase in water content is observed during FOM that could contribute mechanically to the ovulatory process, ultimately leading to oocyte release from surrounding somatic follicular layers (Craik and Harvey, 1984; Milla et al., 2006).

Rainbow trout (Oncorhynchus mykiss) is a major aquaculture species that produces eggs every year starting at age 2 (Bromage et al., 1992). In this species oocyte maturation, ovulation (Cerdà et al., 2007; Grier et al., 2007; Jalabert and Fostier, 1984 ; Lubzens et al., 2010), and egg quality (Bobe, 2015; Bobe and Labbé, 2010; Bromage and Jones, 1992; Brooks et al., 1997) have been extensively studied, including at the molecular level (Bobe et al., 2004, 2006, 2009; Yamashita et al., 2000). As previously documented, exposition to stress factors (e.g., temperature) during FOM can alter the quality of unfertilized eggs released from the ovary at ovulation (Aegerter and Jalabert, 2004; Bobe and Labbé, 2010; Colson et al., 2019).

For these reasons, and because water and ion fluxes are important for several key steps of oocyte development during FOM, the aim of the present study was to investigate the impact of water ionic composition, and more specifically low concentrations of sodium, chloride and potassium on FOM, ovulation and, ultimately, on egg quality, using rainbow trout as a physiological model and relevant aquaculture species. To achieve this objective, we used two closely located production sites in the French Pyrenees where the stream water exhibits dramatically different concentrations of chloride, sodium and potassium all year round, including during the reproductive period.

## 2 Material and methods

### 2.1 Ethical statement

Investigations were conducted in compliance with EU Directive 2010/63/EU for animal experiments. The experimental protocol registered under the reference #201904021747602 was specifically approved by INRAE-LPGP Animal Care and Use Committee. For oocyte sampling, echograph-based monitoring and ovulation detection, fish were anaesthetized in tricaine metanesulfonate (MS222) at a concentration of 130 mg/liter. For gonad sampling, fish were euthanized in MS222 at a concentration of 300mg/liter, until 10 minutes after cessation of opercular movement.

### 2.2 Experimental sites and water characterization

Two experimental sites located in Rébénacq [43.155948,-0.405067] and Sarrance [43.049999,- 0.6014075] (French Pyrenees), and presenting similar water temperature [9°-10°C] and oxygen conditions were used in the present study. The two sites are close (<20 km) but located in different valleys (i.e., hydrogeologic systems) differing mostly by ionic abundance in Sodium (Na+), Chloride (Cl-) and Potassium (K+) in the water (Fig 1.A, Supplemental datafiles 1 and 2). The site presenting significantly lower abundance of these 3 mineral elements will be referred to hereafter as “Low Salinity Water (LSW)”, whereas the other site, for which water ionic composition was significantly higher, will be referred to as “Control”.

**Figure 1:**
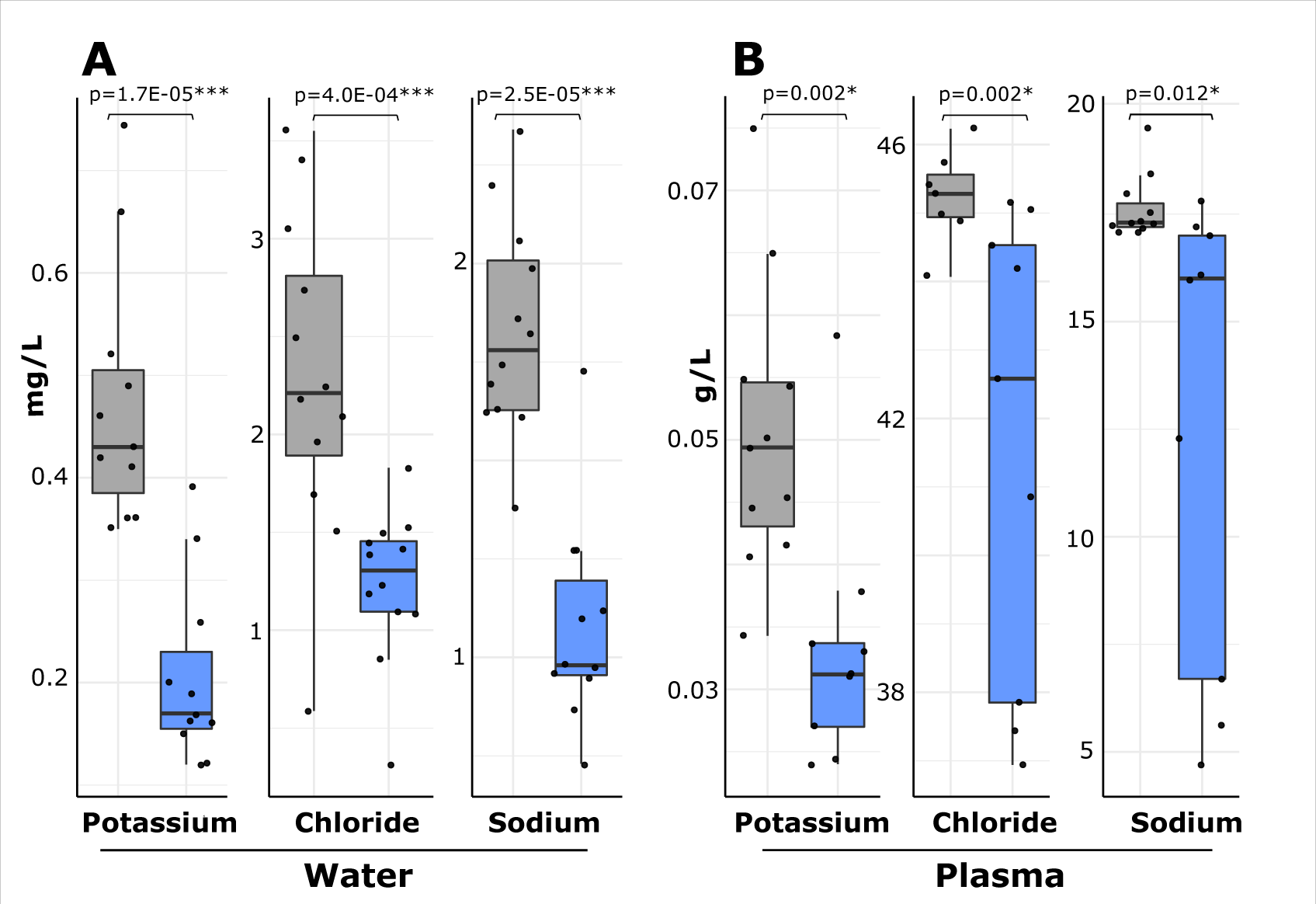
Water and plasma composition on both experimental sites. A: Distribution of Potassium, Chloride and Sodium concentration (mg/L) in Control (in grey, n=12) and Low Salinity Water (in blue, n=12) sites. B: Distribution of Potassium, Chloride and Sodium concentration (g/L) in fish blood plasma reared in Control (in grey, n=11) and Low Salinity Water (in blue, n=9) sites.

For both sites, water composition analyses were performed monthly throughout the year, from the water flowing into the experimental raceways using a dedicated water sampling flask to avoid any contamination. Samples were analyzed by a specialized analytical laboratory (Ultra Traces Analyses Aquitaine, Pau, France) that provided the sampling flasks. Ionic abundance analysis was performed using inductively coupled plasma atomic emission spectroscopy (ICP-AES) technique for Na+ and K+ and by ionic chromatography for Cl-. Data obtained were not normally-distributed (Shapiro-Wilk test, p > 0.005). Ionic concentration in the 2 sites was compared using non parametric U-tests. A comprehensive water analysis of both sites is available in supplemental data files 1 and 2.

### 2.3 Fish rearing

Three years old female rainbow trout (Oncorhynchus mykiss) from the same commercial strain (Viviers SA Company strain, France) were used in this study. After rearing in a nursery site, fish were transferred to either Rébénacq (control site) or Sarrance (LSW site) locations at 6 months post-fertilization. On both sites, fish were reared under natural photoperiod in 125m raceways at a 60kg.m^-3^ density, with a water renewal of 75% per hour until the beginning of the experiment.

Stream water was used in an open circulation system without any recirculation. Similar feeding protocols, including feeding rate and diet composition, were applied for both sites throughout their lifecycle. Fish were fed at an identical daily rate (0.6% of the live weight) with the same commercial diet (46% crude protein, 19% crude fat, 0.9% phosphorus, 11.5% ash, 19 MJ/kg gross energy, Omega- 3 4.8%, Omega-6 1.25%) (Le Gouessant, Lamballe, France). Following a first reproduction at age 2, fish were held under natural photoperiod during their second reproductive cycle. Spawning naturally occurred during November-December.

### 2.4 Experimental design

Approximately two weeks before expected ovulation (estimated based on previous ovulation dates), fish were transferred into a 8 m raceway to facilitate manipulations. Experimental raceways on both sites were of similar dimensions (8 m in length, 1 m in width and 1 m in depth) with a water renewal of 200% per hour originating directly from the stream water. Ovulation was monitored three time a week. Feeding was stopped when first ovulations were detected. On the control site, a total of 32 fish were used including 18 for egg quality assessment using the VisEgg system (Cardona et al. 2020) (4 of them being also used for the oocyte hydration study and 11 of them being also used for blood sampling) and 14 for the molecular analysis. On the LSW site, a total of 46 fish were used including 30 for egg quality assessment using the VisEgg system (4 of them being also used for the oocyte hydration study and 9 of them being also used for blood sampling) and 16 for the molecular analysis.

### 2.5 Egg quality assessment using VisEgg

In both groups, specific fish (control, n=18; LSW, n=30) were individually monitored until ovulation. Ovulations were checked every 2-3 days, under anesthesia, by gentle pressure on the abdomen. Oocyte viability evaluation was systematically performed using the VisEgg protocol when ovulation occurred. This recently developed tool allows a robust assessment of different egg features including egg viability (Cardona et al., 2020). The process uses a standardized image-based batch analysis method of 24h-hydrated unfertilized eggs and an image processing algorithm. Outputs are the presence and percentage of whitening (i.e., non-viable) eggs within the batch. These features can be used to reliably assess egg integrity and viability. Analyze of presence of white eggs depending on rearing sites or the maturation duration yielded contingency tables analyzed using Khi-2 tests. Data obtained from VisEgg were not normally distributed (Shapiro-Wilk test, p < 0.001). Comparison of two groups were performed using non parametric U-tests.

### 2.6 Oocyte development staging during FOM

A non-invasive to the female ultrasound-based staging was used to estimate progress into FOM. This approach, previously described for assessing reproductive stages in wild salmons (Nevoux et al., 2019), was used here for the first time to assess progression into FOM in rainbow trout. An ultrasound scanner M-Turbo (Sonosite) with a 5-10MHz linear transducer (multiple frequency) was used. Anesthetized fish were placed in a plastic tray for a short duration (<30 sec) and sagittal images of the abdominal cavity were obtained by aligning the transducer with the lateral line of the fish. Late vitellogenic oocytes appeared white on the screen, while a dark area progressively appeared at the center of the oocyte as the oocytes progressed into FOM (Fig 3.A).

In order to validate echograph-based staging, direct brightfield observation was performed. In rainbow trout, late-vitellogenic (LV) oocytes observed with the naked eye appear opaque and no noticeable changes in yolk structure or any sign of meiosis resumption can be observed. Post- vitellogenesis (PV) oocytes exhibit a central yolk clarification as meiosis progressively resumes. Germinal vesicle is still visible and lipid droplets remain relatively small. During oocyte maturation (Mat), germinal vesicle breakdown (GVBD) progressively occurs and oocytes become totally clear with well-defined lipid droplets (Grier et al., 2007; Jalabert and Fostier, 1984; Milla et al., 2006; Patiño and Sullivan, 2002; Hurk and Peute, 1979; Wallace and Selman, 1990). The above description of phenotypic changes occurring in the rainbow trout oocyte during FOM was used to validate the echograph-based staging. In 100% of the cases (n= 57 fish) the direct observation confirmed the echograph-based staging of the oocyte during FOM. Echograph-based oocyte staging was preferred due to its non-invasiveness nature and was thus subsequently used to measure the duration of the oocyte maturation phase (i.e., between fully transparent oocyte (Mat) stage and detected ovulation) in the present study. The duration of the maturation phase was temperature-normalized using the actual temperature recorded on both sites during experiments and expressed in hours at 10°C in both experimental groups. Comparisons of the two groups were performed using non-parametric U-tests.

### 2.7 Plasma sampling and analysis

Blood was sampled during oocyte maturation on both sites (control, n=9; LSW, n=11). Blood samples were centrifuged (1000 g, 10 min, 4°C), and plasma were sampled and stored at −20 °C until further analysis. Plasma samples were analyzed by a specialized analytical laboratory (Ultra Traces Analyses Aquitaine, Pau, France). Ionic abundance analysis was performed using inductively coupled plasma atomic emission spectroscopy (ICP-AES) technique for Na+ and K+ and by ionic chromatography for Cl-. Comparison of two groups were performed using non parametric U-tests.

### 2.8 Hydration measurement

For analytical purposes, individual fish originating from the LSW site were separated in two groups, based on the duration of the maturation phase [Mat<64H at 10°C (n=10), Mat 64H at 10°C (n=21)]. Oocyte hydration was measured (n=4 individuals of each group) and monitored from late vitellogenesis stage until ovulation. The method used has previously been described (Milla et al., 2006). Briefly, gentle manual striping was performed under anesthesia to collect oocytes, from late vitellogenesis stage until ovulation. As previously described (Finet et al., 1988), non-ovulated oocytes were expelled from the follicle layers using two fine tip tweezers. Oocyte development staging was performed under a binocular microscope. A total of 60 to 80 oocytes per female were obtained and rapidly weighted to obtain wet mass (WM). After heating these oocytes 24 h at 105°C in a drying oven (Memmert, Buechenbach, Germany), dry mass (DM) was measured. The water content (WC) was measured from subtracting DM from WM. For each individual, WC was measured (n=4 individuals of each group) and monitored from late vitellogenesis stage until ovulation. Analysis of the increase in water content (WC) in the two experimental groups (Mat <64H ; Mat 64H) was performed using non parametric U-test.

### 2.9 Molecular analysis

#### Tissue collection

Ovaries from control (n=4 late-vitellogenesis stage and n=10 oocyte maturation stage) and LSW (n=5 late-vitellogenesis stage and n=11 oocyte maturation stage) groups were sampled for RNA extraction. After euthanasia, ovaries were dissected out of the abdominal cavity. Oocyte staging was performed using ultrasound images prior to euthanasia followed by macroscopic validation of oocyte stage under microscope after dissection. Ovarian follicles were deyolked as previously described (Bobe and Goetz, 2000) by pressing the entire tissue between two stainless steel screens while continuously applying ice-cold Cortland medium with a squirt bottle, then immediately frozen in liquid Nitrogen and stored at -80°C until RNA extraction.

#### Total RNA extraction and reverse transcription

Ovarian tissue was homogenized in Trizol reagent (Molecular Research Center, Cincinnati, USA) at a ratio of 1ml per 100mg of tissue. Tissue was grinded using bead beating technology (Precellys Evolution, Bertin Technologies, Saint-Quentin en Yvelines, France). Total RNA extraction was performed following the manufacturer’s instructions. Briefly, 2 µg of total RNA were reverse transcribed using Maxima First Strand cDNA Synthesis Kit (Maxima Reverse Transcriptase enzyme, derived from Moloney murine Leukemia virus enzyme). Reverse transcription was performed by 10 minutes of incubation at 25°C followed by a 15 minutes step at 60°C and a 5-minute step at 75°C. Control reactions were run without Maxima reverse transcriptase and used as negative controls in the real-time polymerase chain reaction study.

#### Real-time PCR analysis

Real-time PCR was performed using a LightCycler480 (Roche, Switzerland). Reverse transcription products were diluted to 1/30. Technical replicates of each biological sample were run. Runs were performed in 5µL reactions containing 2µL of RT product, 250 nM of each primer and 1X PowerUp SYBR Green Master Mix (Applied Biosystems). The program used was: 50°C for 2 minutes, then 95°C for 2 minutes for initial denaturation, 40 cycles of 95°C for 3 seconds and 60°C for 30 seconds. After amplification, a fusion curve was obtained by 15-second step followed by a 0.5°C increase, repeated 70 times, and starting at 60°C

#### Gene expression analysis

Several concomitant events occur in the ovary during FOM that involve distinct biological processes including ovulation, an inflammatory-like process characterized by major tissue rearrangements, blood coagulation, detachment of somatic follicular cells from the oocyte, and oocyte hydration which involves specific water and ion transports. Genes used for the molecular analysis were thus selected for their putative role in inflammation, proteolytic activity, or water and ion transport and were previously reported to be markedly differentially up-regulated during FOM in rainbow trout (Bobe et al., 2006). Primers used in this prior study (Bobe et al., 2006) were used here to specifically monitor the expression of the following genes. aquaporin 4 (aqp4, GenBank accession # BX885214), vasotocin-neurophysin VT 1 (vt1, GenBank accession # CA375992) and solute carrier family 26 (slc26a4, GenBank accession # BX873066) are involved in ion and water transport processes whereas A Disintegrin And Metalloproteinases 22 (adam22, GenBank accession # CA363158), CXC chemokine L14 (cxcl14, GenBank accession # BX868653) and coagulation factor V (f5, GenBank accession # BX879767) are inflammation-related genes. 18S RNA level was measured in each sample and used for messenger RNA abundance normalization. Expression levels at the late-vitellogenic (LV) stage are not presented but were not significantly different between groups and were arbitrarily set to 1 for each group. Expression data at the oocyte maturation stage were expressed as a percentage of the transcript abundance at the LV stage. Difference between LSW and control groups at oocyte maturation stage were analyzed using non-parametric U-tests.

## 3 Results and discussion

### 3.1 Low water salinity results in lower egg viability

The automatic phenotyping of egg viability using the VisEgg system (Fig 2.A) revealed that 60% of the females exhibited non-viable eggs at the time of ovulation in the LSW group in sharp contrast with the control group in which only 18% of the females exhibited at least one non-viable egg at ovulation. More specifically, the percentage of non-viable eggs was dramatically increased (+378%) in the LSW group in comparison to the control group (Fig 2.B). Salinity is known to have an impact on reproduction in fish, including in salmonids. In Atlantic salmon (Salmo salar) and Coho salmon (Oncorhynchus kisutch), whole-life breeding in freshwater can result in the production of poor quality gametes including poor semen production for males and presence of atretic follicles in females (Valdebenito et al., 2015). Similarly, other studies reported that rearing of Atlantic salmon in sea water induces delays and variability in the timing of ovulation (Haffray, 1995) and poor gamete quality (Maisse et al., 1998). In Arctic charr (Salvelinus alpinus), gamete quality is improved for broodstock transferred in saltwater during their gametogenesis (Atse et al., 2002). Together, these studies indicate that water salinity is a key factor for the reproduction success of salmonids that all reproduce in freshwater even though a part of their life cycle occurs in sea water in some species. Most studies have however focused on the impact of high water salinity during final oocyte maturation. In contrast, data on low salinity water remain scarce. In addition, most studies have focused on migrating species that return to freshwater to reproduce. In the present study, we confirm that water salinity is a key factor of reproductive success and show that, similarly to high salinity, low salinity can also have a detrimental effect on reproductive success.

**Figure 2:**
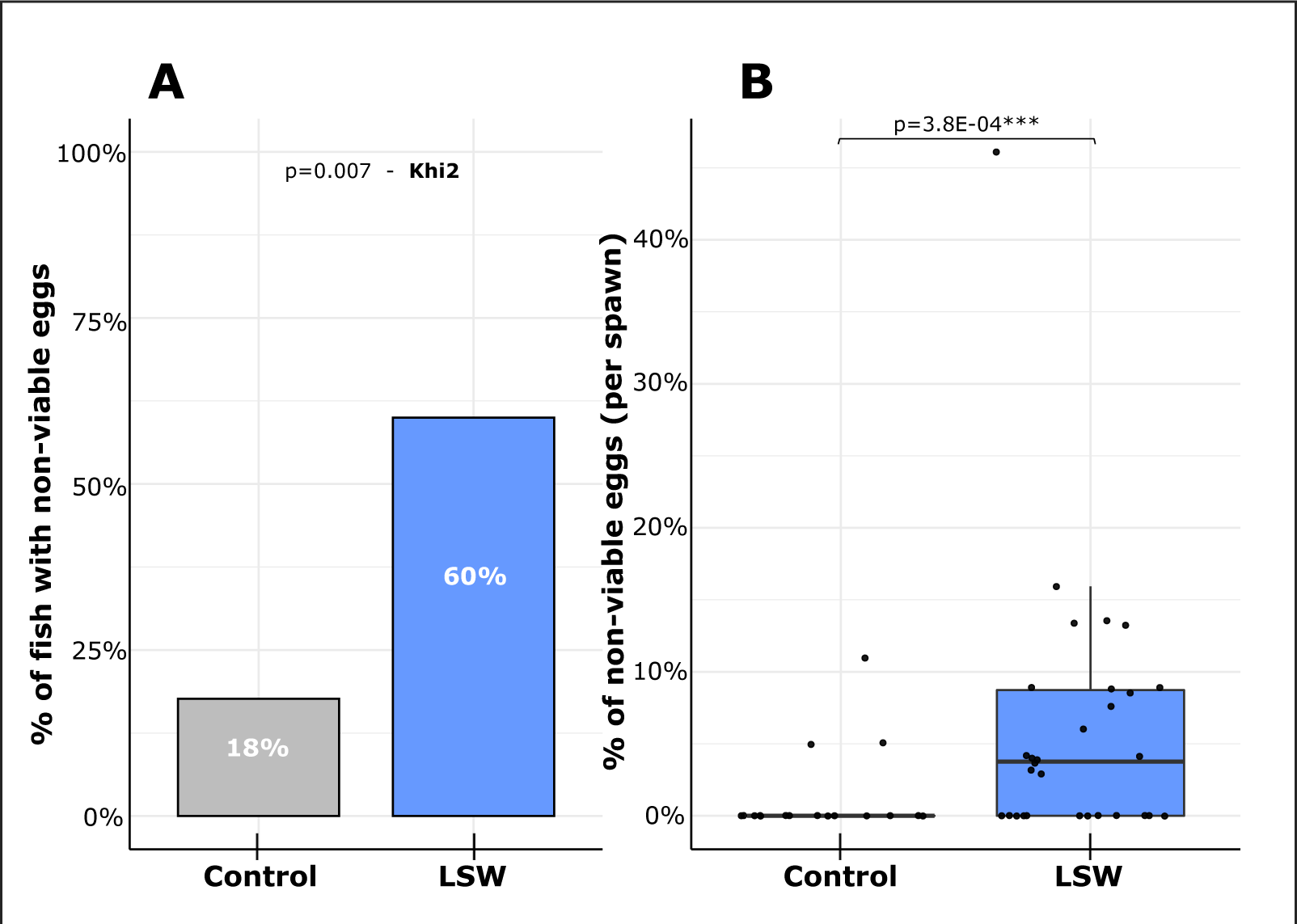
Egg quality in the two experimental sites. A: Occurrence of non-viable eggs in batches evaluated using the VisEgg system in Control (n=18) and LSW (n=30) groups. Khi-2 test (p = 0.007). B: Rate (percent) of non-viable eggs evaluated using the VisEgg system in Control (n=18) and LSW (n=30) groups.

### 3.2 Low water salinity triggers delayed ovulation

Together, our results indicate that low water concentration in chloride, sodium and potassium triggers a significant decrease in egg viability at ovulation. In order to shed light on underlying mechanisms, we aimed at investigating the preovulatory period that is known to be critical for egg quality. Given the favorable temperature range (i.e., 9-10°C) and similar feeding regime throughout the reproductive cycle, we reasoned that LSW could perturbate FOM and investigated the duration of the maturation phase from meiosis resumption to ovulation. Using non-invasive echograph-based analysis we could measure the duration of the oocyte maturation phase. After monitoring the first signs of intra-oocyte clearing (i.e., post-vitellogenic stage) (Fig 3.A and B), oocytes became fully transparent (i.e., maturation stage) until they were released from the ovary at ovulation. The duration of the maturation phase was calculated in each group after normalization by the daily temperature on site. We observed (Fig 3.C) that the maturation phase was significantly longer in the LSW group in comparison to the control group. The duration of oocyte maturation was on average 53 hours (at 10°C) in the LSW group while it was 70 hours (at 10°C) in the control group, which corresponds to a 32% increase in FOM duration in the LSW group.

**Figure 3:**
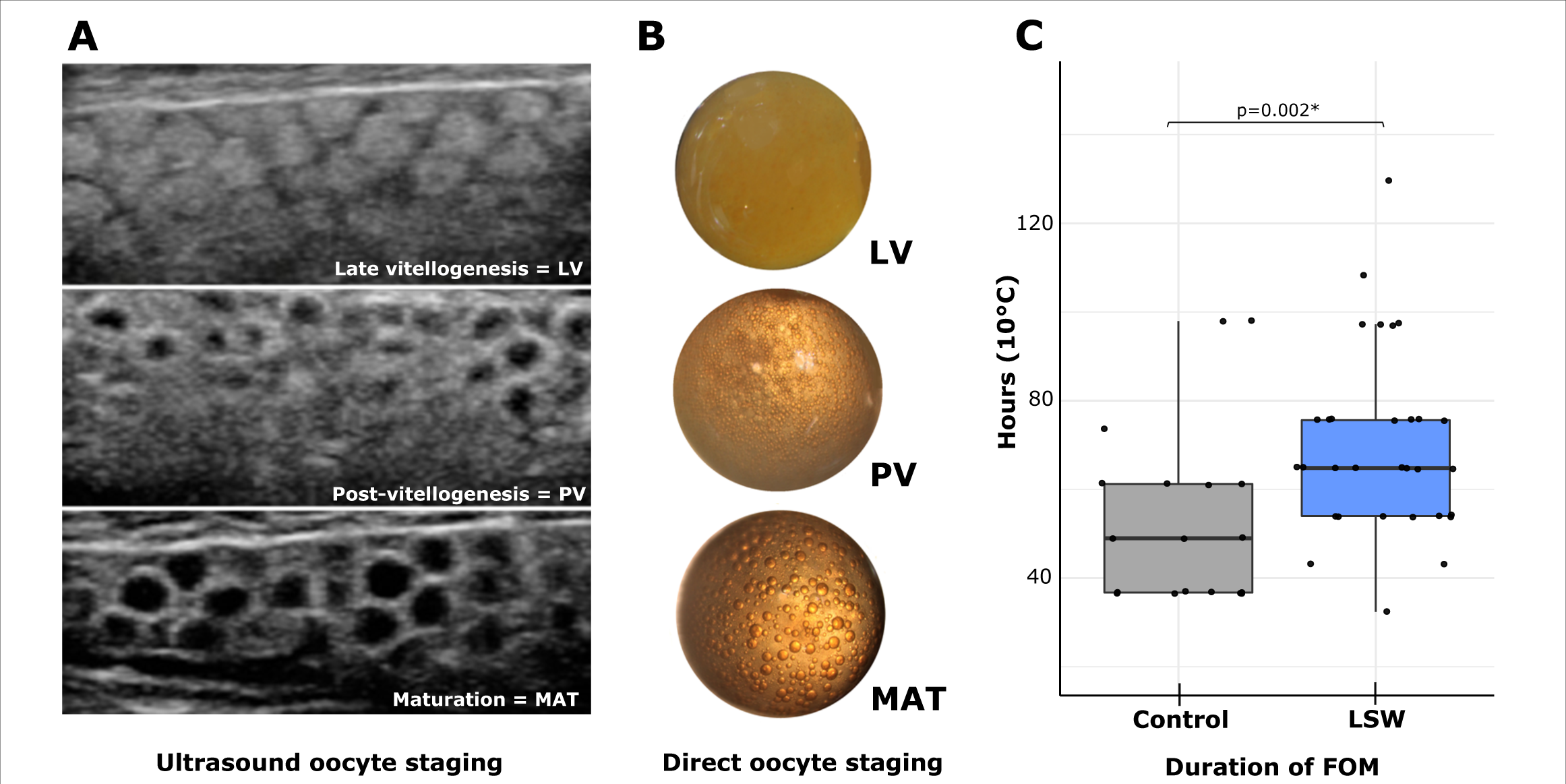
Ultrasound staging for measure of maturation duration. A: Echograph-based observation of oocyte changes during final oocyte maturation. Changes in oocyte aspect from LV to MAT is illustrated by a progressive central darkening, with brightening of peripheral layer before ovulation. B: Pictures of direct observation of oocytes classified by stage. Oocyte were obtained by stripping and defolliculation (Finet et al., 1988). Yolk clarification and lipid droplet formation are observable. C: Distribution of maturation duration evaluated by ultrasound in Control (n=18) and LSW (n=30) from central oocyte clarification and ovulation.

Several studies described the temporal aspects of final oocyte maturation in rainbow trout. The in vivo duration of FOM, including progressive central clearing of the oocyte, ranged from 48 to 72 hours at 13°C (Bry, 1981). The detachment of the mature oocyte from surrounding somatic follicular layers was reported to occur between 72 and 96 hours after the first observable signs of oocyte maturation (i.e., central clarification) at 10°C (Jalabert, 1978; Jalabert and Szöllösi, 1975). Other authors showed that ovulation occurs generally 48-96 hours at 14°C after the peak concentration of maturation-inducing steroid (Fostier, 1981). Another study shown a duration of 48-72 hours at 12°C between maturation and detected ovulation for rainbow trout (Breton et al., 1998).

The duration of oocyte maturation, from full oocyte cytoplasm clearing to ovulation, reported here for the control group, is therefore fully consistent with the duration of FOM reported in these prior studies that usually included the post-vitellogenic phase (i.e., germinal vesicle migration towards the periphery of the oocyte). Our observations however suggest that the duration of FOM in the LSW group is higher or in the upper range of what is classically observed in rainbow trout at 10°C. In addition, the LSW group exhibits a strong interindividual variability with individuals that ovulated 97 hours (2 individuals) and 130 hours (1 individual) after detected oocyte maturation (Fig 3.C). These individuals presented the usual phenotypic traits of maturation (i.e., color pattern and body shape), but ovulation occurred drastically later. Heterogeneity in FOM duration and, in some cases, absence of ovulation has often been reported in trout and other fish. This heterogeneity has however systematically always been associated with premature induction of FOM and ovulation, typically when FOM was triggered in maturationally incompetent (i.e., vitellogenic) follicles in vivo or in vitro. This is for instance the case for hormonally injected fish (Bry, 1981; Goetz and Bergman, 1978; Jalabert, 1978). Using non-invasive ultrasound-based FOM monitoring, we were able to demonstrate here that oocyte maturation (i.e., oocyte clearing) occurred but that ovulation was delayed. Previous studies have demonstrated that oocyte maturation and ovulation, while being interconnected, were two independent processes. In rainbow trout it is possible to trigger ovulation without oocyte maturation (Jalabert, 1972). Together, our results show that low salinity (i.e., low water concentration in chloride, sodium and potassium) during FOM triggers delayed ovulation, sometimes by several days. Our observations also indicate that, while oocyte maturation occurs normally, the ovulatory process appears impaired or inhibited, ultimately leading to abnormally long FOM.

### 3.3 Reduced egg viability in low salinity water is linked with delayed ovulation

Together our results show that low water concentration in chloride, sodium and potassium triggered reduced egg viability and delayed ovulation. We also noticed a high inter-individual variability in both egg viability and maturation phase duration in the LSW group (Figs 2.B & 3.C). We reasoned that delayed ovulation could be at the origin of lower egg viability in the LSW group and investigated further any link between egg viability and maturation phase duration. For data analysis, LSW fish were separated in two groups, based on the duration of the maturation phase. We observed a dramatic and highly significant difference in egg viability in the LSW group depending on the duration of the maturation phase (Fig 4). A very high proportion (85%) of the fish exhibiting a maturation phase duration of 64 hours or more exhibited non-viable eggs at ovulation, while only 10% of the fish that underwent oocyte maturation in less than 64 hours exhibited this feature (Fig 4.A). In addition, the percentage of non-viable eggs was dramatically increased in the 64h group in comparison to the <64h group (Fig 4.B).

**Figure 4:**
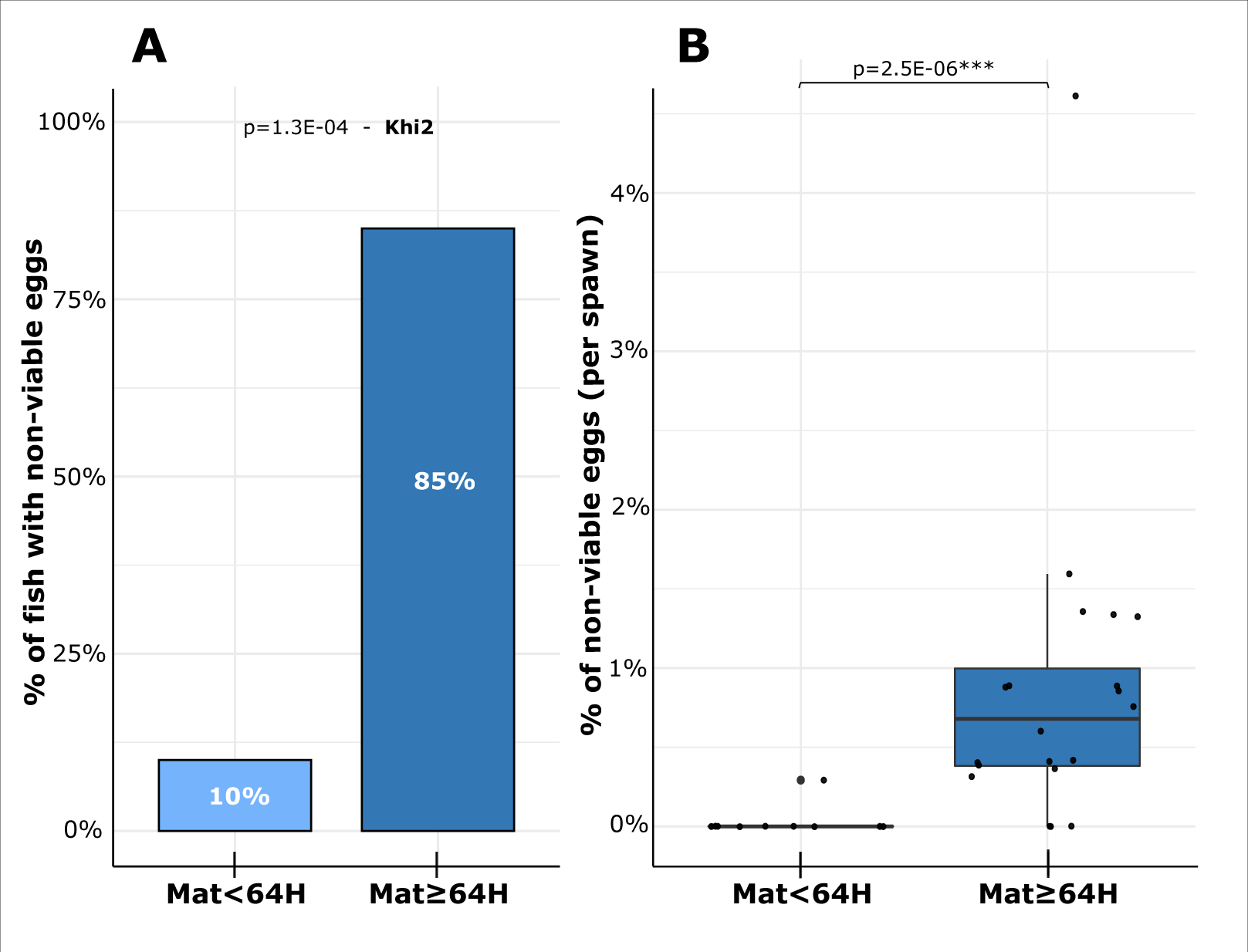
Egg quality depending on the duration of oocyte maturation. A: Percentage of females exhibiting non-viable eggs in low salinity water, evaluated using the VisEgg system. Individuals are separated in two groups depending on the duration of oocyte maturation (Mat<64H, n=10; Mat 64H, n=20). Khi-2 test (p = 1.34E-04). B: Percentage of non-viable eggs depending on the duration of oocyte maturation (Mat<64H, n=10; Mat 64H, n=20).

These results strongly support our hypothesis of a significant loss of egg quality after delayed ovulation. A possible explanation for the higher rate of non-viable eggs would be the ageing of the oocytes. This well-known phenomenon has been extensively described (Aegerter and Jalabert, 2004; Billard, 1981; Craik and Harvey, 1984; Escaffre, 1976; Lahnsteiner, 2000; Samarin et al., 2015, 2008; Springate et al., 1984) and is known to trigger major egg quality defects. All existing studies on oocyte ageing prior to fertilization were however conducted after post-ovulatory ageing. To our knowledge, oocyte ageing prior to ovulation (i.e., within the ovarian follicle) is reported here for the first time in a fish species. In salmonids, in contrast to other fish species, post-ovulatory ageing does not induce a rapid decrease of egg quality. In rainbow trout, ovulated eggs can be held in the body cavity for 5 days at 12°C without any significant decrease of their developmental success (Aegerter et al., 2005). It is therefore unlikely that the modest intra follicular post-meiotic ageing observed here in the LSW group, in which ovulation is delayed, on average, by 17 hours, is sufficient to trigger alone the drop of egg quality observed in LSW. The osmotic properties of the rainbow trout egg have been thoroughly investigated in rainbow trout (Gray, 1932) and another explanation for the increase in whitening egg occurrence could be a loss of the oocyte homeostasis in LSW ultimately leading to non-viable eggs. In brook trout, a diminution of ionic concentration (Cl-) in surrounding medium has as direct negative effect of membrane resistance (Marshall et al., 1985). Disruption of the oocyte homeostasis could therefore originate from a fragile oocyte vitelline membrane. In addition, the loss in oocyte homeostasis could originate from difficulties (i.e., high energetic cost) in maintaining the osmotic difference across oocyte membrane in a hypotonic external medium.

Together, our results show that the reduced egg viability observed in LSW is linked to delayed ovulation. This reduced egg viability can be due, at least in part, to intra-follicular oocyte ageing. We cannot, however, rule out the possibility that LSW also affects egg viability through other mechanisms that could disrupt the maintenance of oocyte homeostasis and ultimately decrease egg viability. Further investigations are needed to unravel the mechanisms triggering decreased egg viability when FOM is occurring in LSW.

### 3.4 Low water concentration in chloride, sodium and potassium results in reduced oocyte hydration during FOM

The release of the oocyte from surrounding follicular layers at the time of ovulation results from the ovulatory process that has originally been described in mammals as an inflammatory-like reaction. In fish, in which the oocyte is directly in contact with surrounding follicular layers, a prerequisite is the separation of the oocyte from surrounding granulosa layers (Jalabert, 1976; Lubzens et al., 2010). This phenomenon has been thoroughly described in rainbow trout in which suppression of tight junctions between granulosa cells and the oocyte results in the apparition of a thin periplasmic space separating the oocyte from surrounding somatic layers (Lubzens et al., 2010). Many observations, including direct brightfield in vitro follicle observation, have shown that preovulatory rainbow trout oocyte exhibits a significant mechanical pression onto surrounding somatic layers (Lubzens et al., 2010). Somatic follicular layers appear to be stretched around the oocyte at that stage. Because of the angiotensin-like nature of the ovulatory process and the contractile actions of theca layers during ovulation (Hajnik and Goetz, 1998; Hsu et al., 1992), postovulatory follicles appear shrinked after ovulation. It was hypothesized that the modest (i.e., 20%), yet significant, increase in water content observed during rainbow trout FOM would contribute to the increasing pressure applied by the oocyte onto surrounding somatic layers and would ultimately mechanically facilitate the exit of the oocyte through a hole in somatic layers (Milla et al., 2006). Mechanic pressure applied by the oocyte swelling is one cause of follicle rupture and the completeness of hydration facilitates the complete liberation of unfertilized eggs into the abdominal cavity. We reasoned that a lower hydration would decrease the efficiency of ovulation and investigated hydration in the two above defined sub-groups exhibiting respectively <64h and 64h maturation phase duration. Oocyte water content (WC) was monitored throughout FOM in both groups. In the <64h group, WC progressively increased throughout FOM in comparison to the late vitellogenic (LV) stage, with a 15% increase during oocyte maturation and a 27% increase at ovulation to reach 36% during the post-ovulatory period (Fig 5, light blue). In contrast, a much more limited increase in water content was observed in the 64h group. In comparison to the LV stage, a 2% increase in water content was observed during oocyte maturation while a 13% increase was monitored at ovulation that did not further increase during the post-ovulatory period. Observations in the <64h group are fully consistent with existing data in rainbow trout that were obtained using the same methodology (Milla et al., 2006). In contrast the hydration in the 64h appears to be drastically reduced. At all stages, the water increase was significantly lower in the 64h group in comparison to the <64h group. Together these results indicate that oocyte hydration is impaired during FOM occurring in low salinity water.

**Figure 5:**
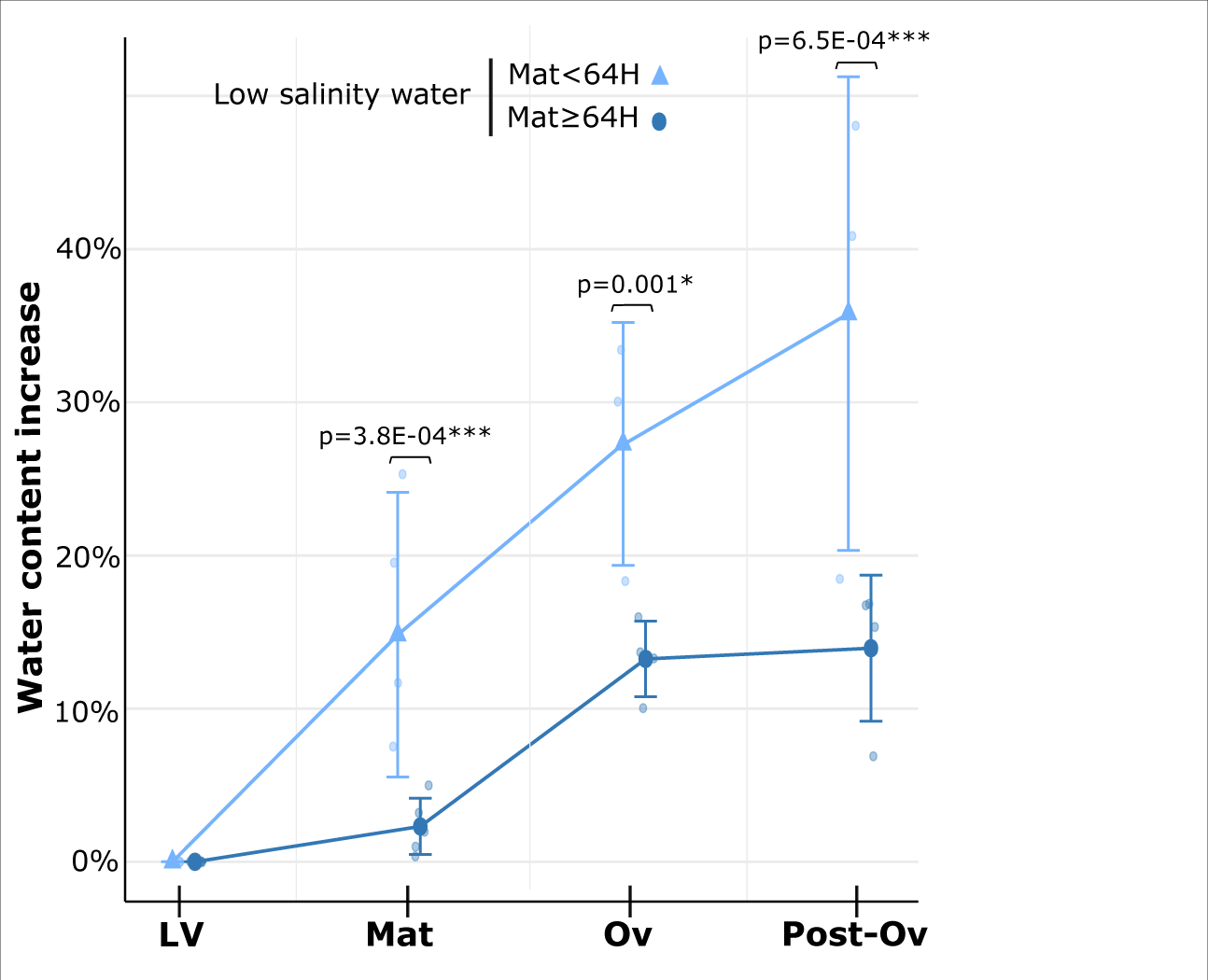
Hydration of oocytes during final oocyte maturation. Increase of oocyte water content (WC), in comparison to late vitellogenic stage, for oocytes sampled at four ovarian stages: Late-Vitellogenesis (LV), oocyte maturation (Mat), ovulation (Ov) and 3 days post-ovulation (Post-Ov). Individuals are separated in two groups based on the duration of oocyte maturation (Mat<64H, n=4; Mat 64H, n=4).

Oocyte hydration has been described for several species (Finn et al., 2002; LaFleur and Thomas, 1991; Milla et al., 2006; Watanabe and Kuo, 1986). Liberation of free amino acids and an uptake of inorganic ions occur during maturation. These events create a change in osmotic equilibrium, resulting in the passive transport of water molecule into the oocyte. This transport mechanism uses several proteins activity such as solute carriers, Na+/K+ ATPases, Aquaporins, and demonstration was made that protein activity and hydration were closely dependent on K+ concentration in the medium (Ecker and Dennis Smith, 1971; LaFleur and Thomas, 1991). K+ and Na+ accumulation in the oocyte during maturation plays an important role in the oocyte hydration process. For Fundulus heteroclitus, oocytes Na+ and K+ uptakes by gap junctions (Lubzens et al., 2010) are main responsible of osmolality increase (Greeley et al., 1991; Wallace et al., 1992). For Atlantic halibut Hippoglossus hippoglossus, Cl- uptakes also contributes to the increase of oocyte osmolality (Finn et al., 2002). Together these studies demonstrate the importance of Na+, K+ and Cl- ions for the oocyte hydration process during FOM. In contrast, the impact of low water salinity was never investigated, mostly because these experiments were often conducted in marine species. We hypothesize that FOM in low salinity water is associated in reduced ions uptake during maturation ultimately leading to incomplete oocyte hydration and thereby causing a large variability in the timing of ovulation. Further investigations are needed to elucidate underlying mechanisms.

### 3.5 Low water concentration in chloride, sodium and potassium induces dysregulated expression of ovulation genes

The ovulatory process involves the activation of a specific gene expression program that has been extensively documented in mammals (Richards et al., 2002) and in fish (Bobe et al., 2009, 2006; Cerdà et al., 2007; Fabra et al, 2005; Hajnik and Goetz, 1998; Knoll-Gellida, 2006). The inflammatory-like nature of the ovulatory process has been well characterized and the expression of many inflammation-related genes have been reported in various fish species (Hagiwara, 2020; Ohnishi, 2005; Takahashi, 2019) including in rainbow trout (Bobe et al., 2006). A subset of these previously identified genes related to inflammation, proteolytic activity, or ion and water transport was monitored to characterize the ovulatory process in low salinity water.

#### Solute transporters and hydration-related genes are under-expressed in low salinity water

Aquaporin are molecular channels responsible of passive water transport across membranes. The role of aquaporins during fish oocyte hydration has been described by several studies. The up- regulation of aquaporin 4 (aqp4) was reported during rainbow trout follicular maturation (Bobe et al., 2006), suggesting that oocyte hydration of fresh water fish is aquaporin mediated, in consistency with the demonstrated role of aquaporin1-like in oocyte hydration of saltwater fish (Cerdà, 2009; Fabra, 2005).

The solute carrier family 26 member 4 gene (slc26a4), previously known as pendrin, codes for a transmembrane anion exchanger and is involved in osmoregulation mechanisms (Nakada et al., 2005). Identified as dramatically up-regulated in the rainbow trout ovary during maturation, slc26a4 is believed to play an important role in the osmotic regulation, causing water influx into oocyte during meiotic maturation. In the present study, a marked under expression of slc26a4 and aqp4 is observed in the ovary when FOM occurs in LSW (Fig 6). Given the role of both genes in ion and water transport, this under expression is fully consistent with the impaired oocyte hydration observed in LSW, in comparison to the control group.

**Figure 6:**
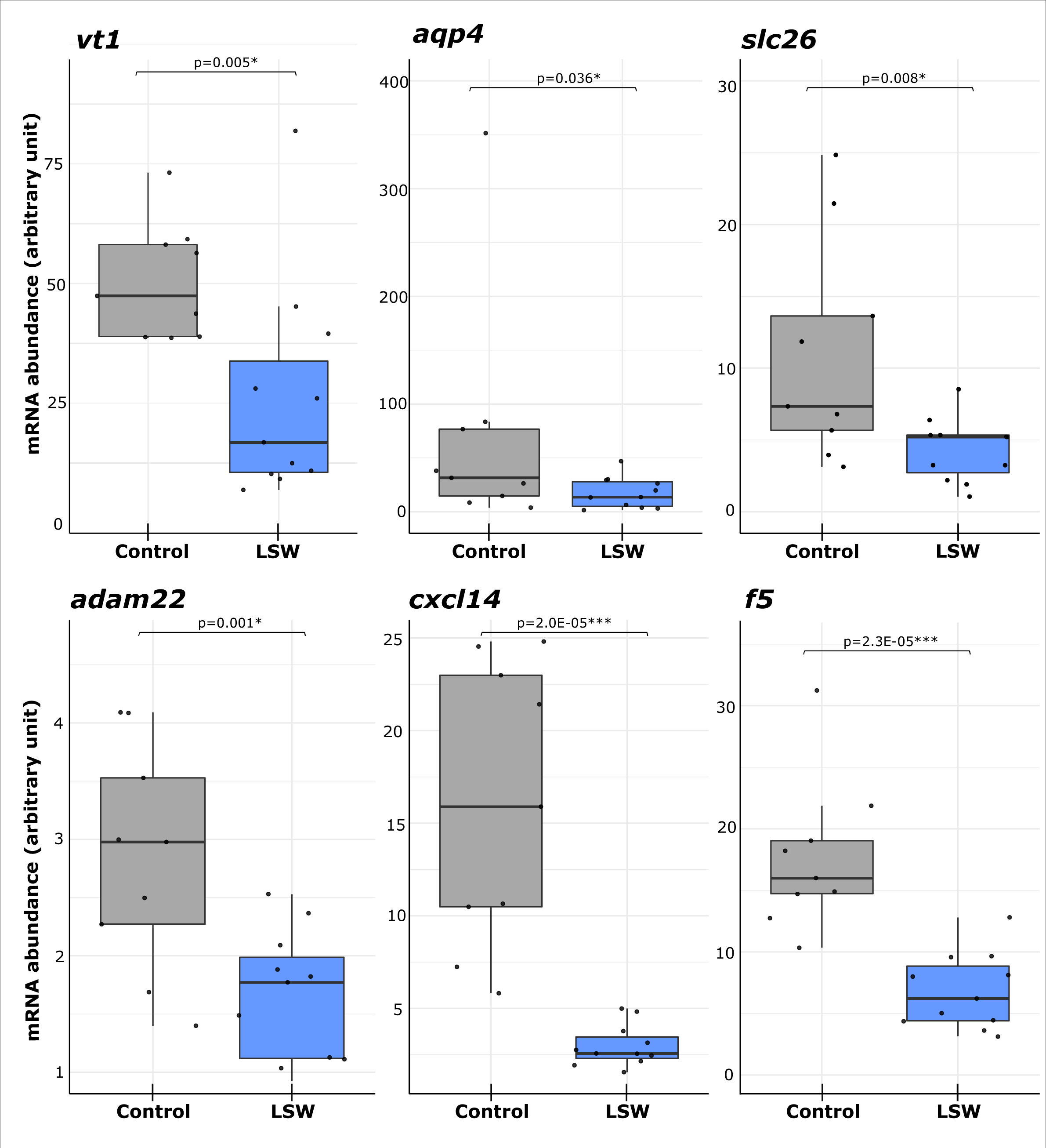
Ovarian expression profiles of genes involved in FOM and ovulation. Ovarian expression profiles of aqp4, slc26, cf5, adam22, cxcl14 and vt5 during oocyte maturation in control (n = 10) and low salinity water (LSW, n = 11) groups. Messenger RNA abundance was measured using real-time PCR and normalized using the abundance of 18S. Abundance was arbitrarily set to 1 in the LV stage in each group.

Vasotocin-neurophysin VT 1 (vt1 gene) has two key roles in FOM-ovulation in catfish (Joy and Chaube, 2015). It is an important link in the gonadotropin cascade controlling FOM and ovulation. By a common action with maturation inductions hormones, VT 1 is a regulator of germinal vesicle breakdown and meiosis resumption. VT 1 also plays a key role in follicular hydration by regulating the aquaporin action (Balment et al., 2006). Here we show a marked under expression of vt1 observed in the LSW group in comparison to the control group (Fig 6). This profile is fully consistent with both the observed impaired ovulatory process and the under expression of aquaporin 4. In addition, the position of vt1 in the gonadotropin-dependent regulatory cascade controlling oocyte maturation and ovulation suggests that it could be a relay in mediating the inhibitory effect of LSW.

#### Inflammation-related genes are under-expressed in low salinity water

Ovulation is accompanied by a broad-spectrum proteolysis within follicular layers that involves a large number of proteases. Members of A Disintegrin and Metalloproteinase gene family are dramatically up-regulated during trout FOM-ovulation phases (Berndtson and Goetz, 1988; 1990; Bobe et al., 2006). Adam22 plays an important role in cell-to-cell and cell-to-matrix interaction regulation (White, 2003). By regulating or processing the deposit of the collagen fibrils, ADAMs proteases cause the extra-cellular matrix proteolysis (Ohnishi et al., 2005).

CXC chemokines stimulate recruitment of leukocytes and play a role of pro-inflammatory mediators. Cxcl14 implication in immune–related functions has been described in fish (Baoprasertkul et al., 2005). The over-expression of cxcl14 during FOM-ovulation phases is fully in line with the inflammatory-like nature of the ovulatory process.

Coagulation factor V (f5 gene) is involved in synthesis of blood coagulant factors and has been characterized in mammals and fish (Hanumanthaiah et al., 2002; Rothberger, 1984). It was speculated that synthesis of coagulation factor V prevents bleeding caused by rupture of follicle layers during ovulation.

Here we show that cxcl14, adam22, and f5 are dramatically under expressed during oocyte maturation in the LSW group in comparison to the control group (Fig 6). These expression profiles are consistent with the delayed ovulation observed in LSW and suggest that the inflammatory nature of ovulation is affected. This is especially true for cxcl14 for which a 5-fold decrease in gene expression is observed in the LSW group in comparison to the control group. Together, these results show that several important components of the ovulatory process are toned down when FOM occurs in low salinity water. It is noteworthy that most, if not all, aspects of the ovulatory process including inflammation, coagulation, swelling, water and ion transport as well and proteolysis are under activated. Even if previous studies have shown gene expression changes when fish adapt to water salinity change (e.g. migration, precipitation) for salmonids (Morro, 2020; Norman et al., 2014) and other species (Fiol, 2006; Pujante, 2018), evidence of such an impact of low water salinity on the overall ovulatory process were, to our knowledge, previously unreported. This raises the question of how low water salinity can inhibit the overall ovulatory process and not only osmotic exchanges.

### 3.6 Lower blood plasma levels of sodium, chloride and potassium are observed in the LSW site

In order to assess the impact of water concentration in chloride, sodium and potassium on fish homeostasis, we measured blood plasma concentration in these three elements in both control and LSW sites. We observed a significantly lower plasmatic concentration of these three mineral elements in fish of the LSW site in comparison to the control site (Fig 1.B). This indicates that the water composition can lead to significant differences in plasma concentration in these elements. This also further supports our hypothesis that reproduction in fresh water exhibiting low sodium, chloride and potassium concentration perturbates the fish oocyte osmolarity and the hydration process during final oocyte maturation and ovulation.

### 3.7 Role of other water components

While marked and highly significant differences in water concentration of chloride, sodium and potassium are found between the two experimental sites, other differences exist in water composition that could also contribute, at least in part, to the impaired FOM and ovulation and reduced egg quality phenotype observed in the LSW site. A lower concentration in Iron (Fe) and Manganese (Mn) is observed in the LSW site in comparison to the control site. For Iron, the difference is however not significant when only the reproductive period is considered (Supplemental datafile 2). Manganese (Mn) is known to inhibit ovulation in rainbow trout (Jalabert and Szollosi, 1975) and inhibit meiotic maturation in terrestrial animals (Bilodeau-Goeseels, 2001). It is therefore difficult to link low Mn concentration with impaired FOM and ovulation. In addition, it is commonly accepted that the diet is considered to be the major source of Manganese and Iron (Lall, 2002). It is therefore extremely unlikely that the lower water concentration in Mn and Fe found in the LSW site is responsible for the impaired reproductive phenotype observed in the LSW site. Similarly, a lower nitrate concentration was observed in the LSW site. While we cannot rule out that this could contribute to the abnormal reproductive phenotype observed in the LSW site, this seems however, to our knowledge and in the light of previous studies led on salmonids and others fish species (Dolomatov, 2011; Tripathi and Krishna, 2008), extremely unlikely.

## 4 Conclusion

Here we show that low water concentration in chloride, sodium and potassium negatively influences final oocyte maturation, ovulation and, ultimately, egg quality. Low salinity water triggers delayed ovulation and lower oocyte viability. When investigating the impact of LSW on FOM duration, individuals presenting the most severe phenotypes exhibited impaired oocyte hydration and abnormally reduced gene expression levels of several key players of the ovulatory process. While the under expression of water (i.e., aquaporins) and ion (i.e., solute carriers) transporters is consistent with impaired oocyte hydration, our observations also indicate that the entire ovulatory gene expression sequence is disrupted. Our results raise the question of the mechanisms underlying the negative influence of low water concentration in chloride, sodium and potassium on the dynamics of the preovulatory phase, on the control of the oocyte homeostasis, including hydration, and on the overall success of the maturation-ovulatory process. A global analysis of differentially regulated genes in LSW during final oocyte maturation is needed to unravel these mechanisms.

## 5 Declarations

### 5.2 Authors’ contributions

ES conceived the study, performed experiments and data analysis and drafted the manuscript. EC participated in experiments. SS-K and FC participated in the design of the study. JB conceived the study and participated in manuscript writing and data analysis.

## Acknowledgements

The authors thank Floriane Colon and Viviers Company staff for their help in fish rearing. The authors thank the Nouvelle-Aquitaine regional council and ANRT for the financial support which allowed the study.

## >7 Additional Files

**Supplemental data file 1:** Water composition in control (Rébénacq) and LSW (Sarrance) sites. Mean (N= 12) yearly (2020-2021) values and standard deviation are shown. Difference between LSW and control water were analyzed using non-parametric U-tests. * denotes significant differences.

7 Additional Files
|  | CONTROL |  | LSW |  |
| --- | --- | --- | --- | --- |
|  | Mean | SD | Mean | SD |
| Conductivity ($\mu\text{S}/\text{cm}$) | **264,00** | 41,64 | **248,92** | 34,07 |
| **Major ions, in milligrams per liter** |  |  |  |  |
| Bicarbonate | **145,50** | 17,77 | **140,00** | 17,04 |
| Calcium | **44,79** | 8,57 | **43,60** | 5,78 |
| Chloride | **2,29** | 0,84 | **1,24** | 0,39 |
| Magnesium | **4,29** | 0,51 | **5,16** | 2,41 |
| Potassium | **0,47** | 0,13 | **0,21** | 0,09 |
| Sodium | **1,83** | 0,28 | **1,09** | 0,27 |
| Sulfur | **3,03** | 0,79 | **5,08** | 3,32 |
| Sulfate | **8,33** | 2,06 | **13,05** | 9,54 |
| **Nutrients, in milligrams per liter** |  |  |  |  |
| Nitrate | **2,92** | 1,28 | **1,03** | 0,27 |
| **Trace elements, in micrograms per liter** |  |  |  |  |
| Copper | **0,22** | 0,09 | **0,14** | 0,07 |
| Iron | **14,99** | 14,12 | **3,00** | 4,58 |
| Manganese | **1,47** | 1,28 | **0,15** | 0,18 |
| Selenium | **0,13** | 0,02 | **0,15** | 0,05 |
| Iode | **1,35** | 0,53 | **1,04** | 0,18 |
| Zinc | **1,37** | 0,70 | **1,35** | 0,83 |

\*
\*
\*
\*
\*

**Supplemental data file 2:** Water composition in control (Rébénacq) and LSW (Sarrance) site during the reproductive period (October-December). Mean (N= 3) values (from 2020-10-15 to 2020-12-15) and standard deviation are shown. Difference between LSW and control water were analyzed using non-parametric U-tests. * denotes significant differences.

|  | CONTROL |  | LSW |  |
| --- | --- | --- | --- | --- |
|  | Mean | SD | Mean | SD |
| Conductivity (µS/cm) | **282.00** | 20.785 | **242.67** | 29.48 |
| **Major ions, in milligrams per liter** |  |  |  |  |
| Bicarbonate | **154.50** | 25.05 | **143.67** | 10.97 |
| Calcium | **48.97** | 5.11 | **42.43** | 4.66 |
| Chloride | **2.98** | 0.45 | **1.47** | 0.06 |
| Magnesium | **4.59** | 0.32 | **3.51** | 2.66 |
| Potassium | **0.60** | 0.19 | **0.18** | 0.04 |
| Sodium | **2.02** | 0.17 | **0.99** | 0.18 |
| Sulfur | **3.60** | 0.09 | **3.10** | 2.82 |
| Sulfate | **9.70** | 0.65 | **9.5** | 8.37 |
| **Nutrients, in milligrams per liter** |  |  |  |  |
| Nitrate | **4.09** | 1.79 | **1.11** | 0.09 |
| **Trace elements, in micrograms per liter** |  |  |  |  |
| Copper | **0.26** | 0.15 | **0.16** | 0.13 |
| Iron | **8.07** | 6.72 | **2.00** | 0.44 |
| Manganese | **0.70** | 0.33 | **0.12** | 0.07 |
| Selenium | **0.13** | 0.01 | **0.17** | - |
| Iode | **1.45** | 0.50 | **1.15** | 0.08 |
| Zinc | **1.26** | 0.56 | **1.44** | 0.54 |

\*
\*
\*
\*
\*

